# *In situ* quantification of osmotic pressure within living embryonic tissues

**DOI:** 10.1101/2022.12.04.519060

**Authors:** Antoine Vian, Marie Pochitaloff, Shuo-Ting Yen, Sangwoo Kim, Jennifer Pollock, Yucen Liu, Ellen Sletten, Otger Campàs

## Abstract

Mechanics is known to play a fundamental role in many cellular and developmental processes. Beyond active forces and material properties, osmotic pressure is believed to control essential cell and tissue characteristics. However, it remains very challenging to perform *in situ* and *in vivo* measurements of osmotic pressure. Here we introduce double-emulsion droplet sensors that enable local measurements of osmotic pressure intra- and extra-cellularly within 3D multicellular systems, including living tissues. After generating and calibrating the sensors, we measured the osmotic pressure in blastomeres of early zebrafish embryos as well as in the interstitial fluid between the cells of the blastula by monitoring the size of droplets previously inserted in the embryo. Our results show a balance between intracellular and interstitial osmotic pressures, with values of approximately 0.7 MPa, but a large pressure imbalance between the inside and outside of the embryo. The ability to measure osmotic pressure in 3D multicellular systems (developing embryos, organoids, etc.) will help understand its role in fundamental biological processes.

---

Mechanics has been shown to affect fundamental biological processes across scales, from cellular function to organ formation and tissue homeostasis^1–5^. Actomyosin force generation, cell-cell adhesion, traction forces and membrane tension have all been shown to affect cellular activity at subcellular and cellular scales^5^. At a multicellular level, active force generation^2,6,7^ and spatiotemporal control of tissue material properties^8,9^ have been shown to play a key role in tissue morphogenesis during embryonic development, as well as in the control of cell migration^10^ and cell differentiation^11,12^. Other fundamental cellular and developmental processes, such as the control of cell and nuclear sizes^13–15^, cell division^16^, cytoskeletal mechanics^17,18^, the emergence of a blastocoel in early mammalian embryos^19^, the formation of complex lumen structures during organogenesis (liver, pancreas, lung, etc.)^20–22^, and the emergence of gradients in extracellular spaces during embryonic development^9,23^, all depend on a tight control of the osmotic pressure both inside cells and in the extracellular space^24,25^. Yet, measuring osmotic pressure remains very challenging, especially in 3D multicellular systems such as living tissues or organoids, hindering our understanding of the role that osmotic pressure plays in living organisms.

Previous measurements of the hydrostatic pressure difference across the cell surface in animal cells *in vitro*, or in externally accessible lumens *in vivo*, have been achieved using either microneedles as a pressure gauge or other surface contact probes, such as atomic force microscopy^16,26–28^. These techniques require an external probe to be in constant contact with the sample, which is invasive and not well-suited for 3D multicellular systems that continuously change shape. Intracellular osmotic pressures in animal, fungal and plant cells have been estimated *in vitro* by applying osmotic shocks, with estimated values ranging between 0.1-1 MPa^17,18,29^. Previous microdroplet-based techniques have been developed to measure mechanical stresses^30^ or material properties^31^ *in situ* and *in vivo*, but these do not allow measurements of (osmotic) pressure. Gel beads can perform measurements of isotropic stress associated with cellular crowding in multicellular systems, but cannot measure osmotic pressure either^32,33^. Finally, measurements of the interstitial fluid osmolarity in early zebrafish embryos were achieved using standard osmometers by collecting large interstitial fluid quantities in whole tissue explants^34^. These measurements provided an average value of interstitial fluid osmolarity for the entire explant, which required the destruction of the sample, thereby precluding any measurements of spatial or temporal variations in osmotic pressure in the tissue. Measuring osmotic pressure locally *in situ* and *in vivo*, within cells or tissues of developing embryos (including lumen formation in organogenesis), or in other 3D multicellular systems such as organoids, remains challenging.

Here we introduce novel osmotic pressure sensors able to quantify osmotic pressure intra- and extra-cellularly within 3D living tissues, including developing embryos. The sensors consist of double emulsion microdroplets made of a biocompatible oil droplet containing a smaller aqueous droplet with a calibrated concentration of osmolyte (Fig. 1a). The oil surrounding the inner aqueous droplet acts as a protective shell while simultaneously allowing surfactant-mediated water transport, effectively behaving as a water permeable layer (Fig. 1a). By controlling the osmolyte concentration in the inner droplet, as well as the surfactants in the oil and the relative inner/outer droplet sizes, it is possible to generate osmotic pressure sensors with well-defined characteristics. After calibrating the sensors, we used them to measure the osmotic pressure in blastomeres (cells) of early zebrafish embryos, as well as in the interstitial fluid between the cells of the zebrafish blastula. Our results show that double-emulsion droplets enable *in situ* and *in vivo* measurements of osmotic pressure, both intra- and extra-cellularly within living embryonic tissues.

**Figure 1.**
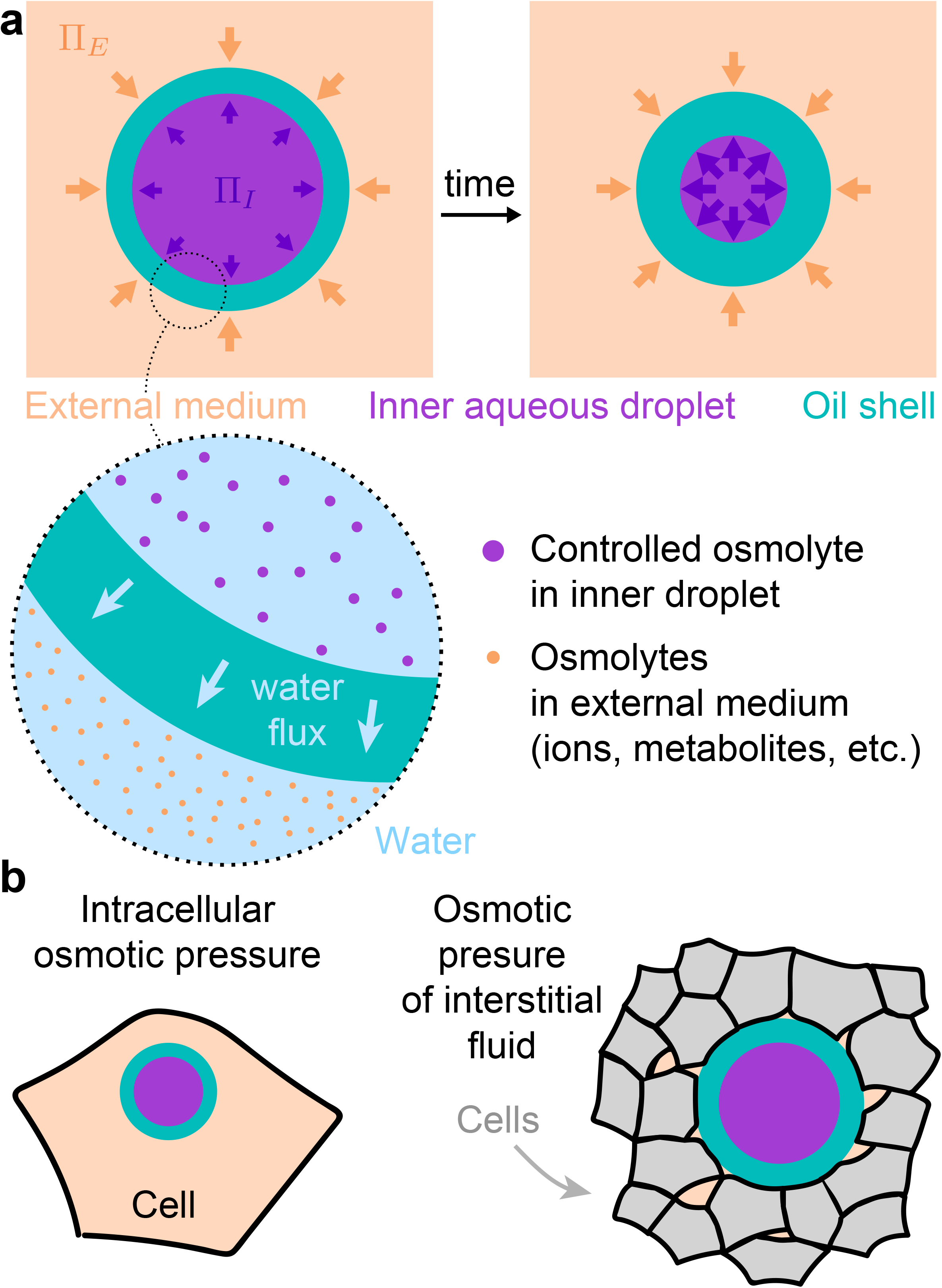
Double-emulsion droplets as osmotic pressure sensors. **a,** Sketch of double-emulsion droplets used as osmotic pressure sensors in cells or in the interstitial space between cells withing living tissues. Relevant physical parameters are defined. **b,** Sketch of a double emulsion droplet inside a cell (left) and in the extracellular space between cells (right), enabling measurements of the intracellular osmotic pressure and of the osmotic pressure of the extracellular interstitial fluid, respectively.

## Double-emulsion microdroplets as osmotic pressure sensors

Double emulsion droplets, composed of an aqueous droplet embedded in an oil shell (water-in-oil-in-water, or W/O/W double emulsions), could potentially be used as osmometers since water flux through the oil shell is possible in the presence of surfactants^35–37^ (Fig. 1a). Such water transport is thought to rely on inverse micelles formed by the surfactant within the oil layer^38–41^. Thanks to the outer water-permeable oil layer, the inner aqueous droplet can increase or decrease its volume as water enters or leaves the droplet, respectively (Fig. 1a). Previous studies have shown that changes in osmolarity in the external medium can drive water flows through the oil shell of the double-emulsion droplet^35,37,38,42^, indicating that the system is sensitive to osmotic pressure differences. In order for double-emulsion droplets to be used as osmotic pressure sensors, the osmolarity and size of the inner aqueous droplet, as well as the size and surfactant composition of the outer oil layer must be controlled, enabling the generation of stable and calibrated double-emulsion droplets.

To produce monodispersed, stable water-in-oil-in-water double-emulsion droplets we used droplet microfluidics^43^ (Fig. 2a-c; Methods), as it enables control over the initial volumes of both the inner aqueous droplet and outer oil layer within our desired range (10-40 μm in outer droplet radius). To ensure biocompatibility, we used fluorocarbon oils for the oil phase and non-ionic fluorinated surfactants (Krytox-PEG) to stabilize the droplets (Methods), which have both previously been extensively used in biological applications^44^, including as *in vivo* mechanical stress sensors and actuators^9,30,31,45^. In addition, we used a fluorinated fluorophore^46^ to visualize the oil layer using fluorescence microscopy (Methods). Finally, in order to control the osmotic pressure of the inner aqueous droplet, we introduced high molecular weight polyethylenglycol (PEG; a small fraction it being fluorescently-labeled) at a controlled concentration as a water soluble non-ionic osmolyte during the generation of droplets (Fig. 2a,b; Methods). Microfluidic generation of such droplets in a polyvinyl alcohol (PVA) solution of fixed osmolarity led to stable double emulsions with controlled initial volumes (Fig. 2c; Methods). The fluorescent dyes in the inner droplet and in the oil layer enable the quantification of inner/outer droplet sizes at high resolution using fluorescence confocal microscopy (Fig. 2d).

**Figure 2.**
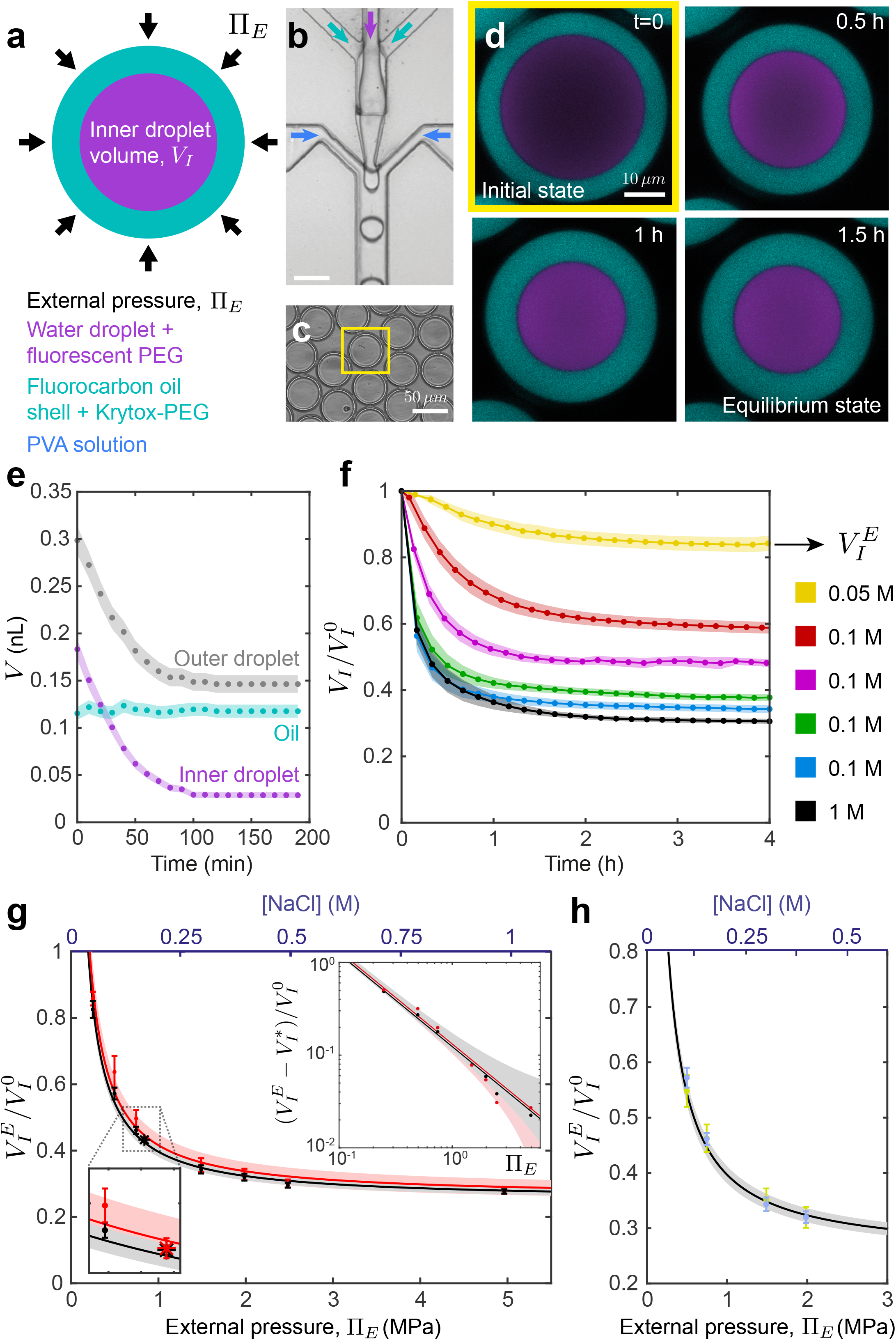
Characterization of double-emulsion droplets at equilibrium. **a,** Sketch of a double-emulsion droplet indicating its composition and characteristics. **b-c,** Microfluidic generation (**b**) of double-emulsion droplets (**c**). Scale bar, 50 μm. **d,** Confocal section of a double-emulsion droplet showing the temporal reduction in inner and outer droplet sizes when placed in a 0.4 M NaCl solution. Fluorocarbon oil (cyan) and fluorescent PEG (purple) are shown (color code as sketched in **a**). Scale, 10 μm. **e,** Temporal evolution of the inner droplet volume, *V_t_* (purple), the outer droplet volume, *V_T_* (gray) and the oil layer volume (cyan). Error bands are measurement error for single droplet. **f,** Temporal evolution of the inner droplet volume, *V_I_* (normalized by the initial volume 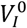 of the inner droplet at equilibrium with a 10% w/w PVA solution; 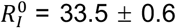, with 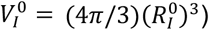), for double-emulsion droplets with fixed initial internal PEG concentration placed in salt solutions of varying osmolarities (Methods). Error bands are SD. N=20 (yellow), 16 (red), 16 (purple), 15 (green), 21 (blue), 17 (black). **g,** Measured dependence of the equilibrium inner droplet volume, 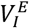 (normalized by 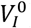), on the externally imposed osmolarity (or osmotic pressure, Π_*E*_) for droplets with 5% w/w (black circles) and 10% w/w (red circles) initial PEG concentrations. The top inset shows the power-law dependence of the normalized osmotically-active equilibrium volume of the inner droplet, 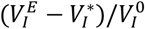, on the external osmotic pressure, Π_*E*_, for both PEG initial concentrations. Black and red lines are fits of Eq.1 to the data for each initial PEG concentration. The small inset shows a magnified region of **g** with the measured equilibrium volumes of the inner droplet of double-emulsion droplets with 5% w/w (black star) and 10% w/w (red star) initial PEG concentrations when placed in cell culture media of known osmolarity (Methods). N=13 (black), 25 (red). Fit CBs (68%) are shown. Error bars are SD. **h,** Measured dependence of the equilibrium inner droplet volume, 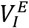, on the externally imposed osmolarity (or osmotic pressure, Π__E__) for droplets of different initial sizes (large droplets: 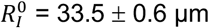, blue; small droplets: 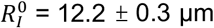, green) but same initial PEG concentration (5% w/w). Black line is the calibration curve (fit in **g**) for 5% w/w PEG. Small droplets: N=26 (0.5 MPa), 23 (0.75 MPa), 18 (1.5 MPa), 23 (2 MPa). Large droplets: N=16 (0.5 MPa), 16 (0.75 MPa), 15 (1.5 MPa), 22 (2 MPa). CB (68%) of calibration curve is shown. Error bars are SD.

Once produced, we characterized the response of double-emulsion droplets to controlled changes in osmolarity in the external medium. Placing double-emulsion droplets in an aqueous medium containing a salt (NaCl) concentration of 0.4M drove a progressive and strong reduction in droplet volume as water left the inner droplet through the oil layer (Fig. 2d). This led to an increase in the fluorescent intensity signal in the inner droplet as fluorescent PEG became more concentrated. Monitoring the reduction in the volumes of both inner and outer droplets (Fig. 2a; Methods) showed that both decreased equally over time from their respective initial volumes, eventually reaching equilibrium volumes as the pressure in the inner droplet equilibrated with the external pressure (Fig. 2e). Throughout this process, the oil shell volume remained constant (Fig. 2e), as expected for fluorocarbon oils with low water solubility^38,47^, indicating that monitoring the inner or outer droplet volume provides the same information about droplet sizes. The fact that the reduction in volume reached an equilibrium value indicates that only water is transported through the oil shell. Indeed, long term (12h) imaging of double-emulsion droplets at varying laser intensities displayed a laser power dependent decay in fluorescent PEG signal intensity, indicating that the slight observed decay is mostly due to photobleaching rather than PEG leakage from the inner droplet (Supplementary Fig. 1). These results indicate that double-emulsion droplets have the necessary characteristics to be used as proper osmotic pressure sensors.

To test the sensitivity of double-emulsion droplets to different external osmotic pressures, we monitored the temporal evolution of their inner droplet volume *V_I_* when placed in salt (NaCl) solutions of different concentrations, ranging from 0.05M to 1M (Fig. 2f). The osmolality of each of these solutions was measured using a commercial osmometer, allowing us to obtain the osmotic pressure Π_*E*_ of each solution (ranging from 0.25 to 4.96 MPa; Methods; Supplementary Fig. 2). These salt solutions of known osmolalities (and osmotic pressures) were used to calibrate the double emulsion droplets. For all concentrations, the inner droplet volume decreased over time from its initial volume 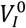 until reaching an equilibrium volume 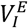 that depended on the externally imposed osmotic pressure Π_*E*_, with larger osmotic pressures leading to smaller equilibrium volumes (Fig. 2f). The equilibrium volume of the inner droplet showed a power law dependence on the external pressure (Fig. 2g and top inset), albeit never becoming smaller than a minimal volume, 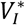, associated with PEG volume exclusion (osmotically inactive volume), as previously reported^17,18^. This power law relation is consistent with the inner droplet’s osmotic pressure 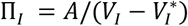 being equal to the external osmotic pressure at equilibrium, namely

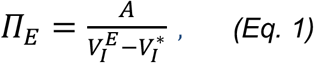

where *A* is a constant associated with the inner droplet osmolyte concentration and can be related to the initial conditions of droplet preparation by 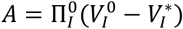, with 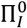 being the osmotic pressure of the initial PVA solution, fixed in our experiments at 79 mOsm/kg or 0.2 MPa (Methods). Double-emulsion droplets with different initial PEG concentrations in the inner droplet also follow Eq. 1 (Fig. 2g). To test if this same relation holds in the presence of more complex external chemical environments, we placed double-emulsion droplets in cell culture media (Methods). The resulting equilibrium inner droplet volumes follow the same relation in cell culture media as for simple salt solutions with the same osmotic pressure, regardless of the initial PEG concentrations in the inner droplet (Fig. 2g, small inset). Finally, the same behavior was also observed for different initial inner droplet volumes at fixed PEG concentration in the inner droplet (Fig. 2h). These results indicate that the power law relation in Eq. 1 constitutes a robust calibration curve of double-emulsion droplets, providing the relation between the measured inner droplet volume and the osmotic pressure in the external medium at equilibrium.

Beyond equilibrium values, to evaluate the temporal resolution of the measurements, it is important to know the relaxation timescale *τ_R_* of pressure equilibration in double-emulsion droplets. To that end, we monitored the volume of the inner droplet over time and measured the dependence of the relaxation timescale on the different control parameters (Fig. 3a-b; Methods), namely the initial inner droplet radius, 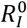, initial internal pressure, 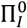, imposed external pressure, Π_*E*_, and the initial oil volume fraction, 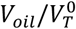 (Fig. 3a; Methods). The relaxation timescale *τ_R_* displayed a strong dependence on the initial inner droplet size 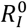, with increasing relaxation time for increasing droplet sizes (Fig. 3c). While smaller values of the initial inner pressure 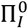 led to shorter relaxation timescales (Fig. 3d), pressure equilibration occurred faster for larger external pressures Π_*E*_ (Fig. 3e). Finally, no dependence of the relaxation timescale on the oil volume fraction 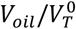 was observed (Fig. 3f), likely because there is always a region where the oil layer thickness is small due to the inner droplet buoyancy (Fig. 3f, inset). The measured values of *τ_R_* were not affected by the presence of fluorinated dye in the fluorocarbon oil (Supplementary Fig. 3). In order to perform measurements of osmotic pressure on relatively short timescales (~10 min), the initial radius of the inner droplet 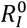 should be smaller than approximately 20 μm and have small initial internal pressures (<100 kPa; Fig. 3c). In what follows, we generate droplets with these characteristics (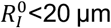 and 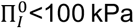; Fig. 3c) to perform osmotic pressure measurements in living embryos.

**Figure 3.**
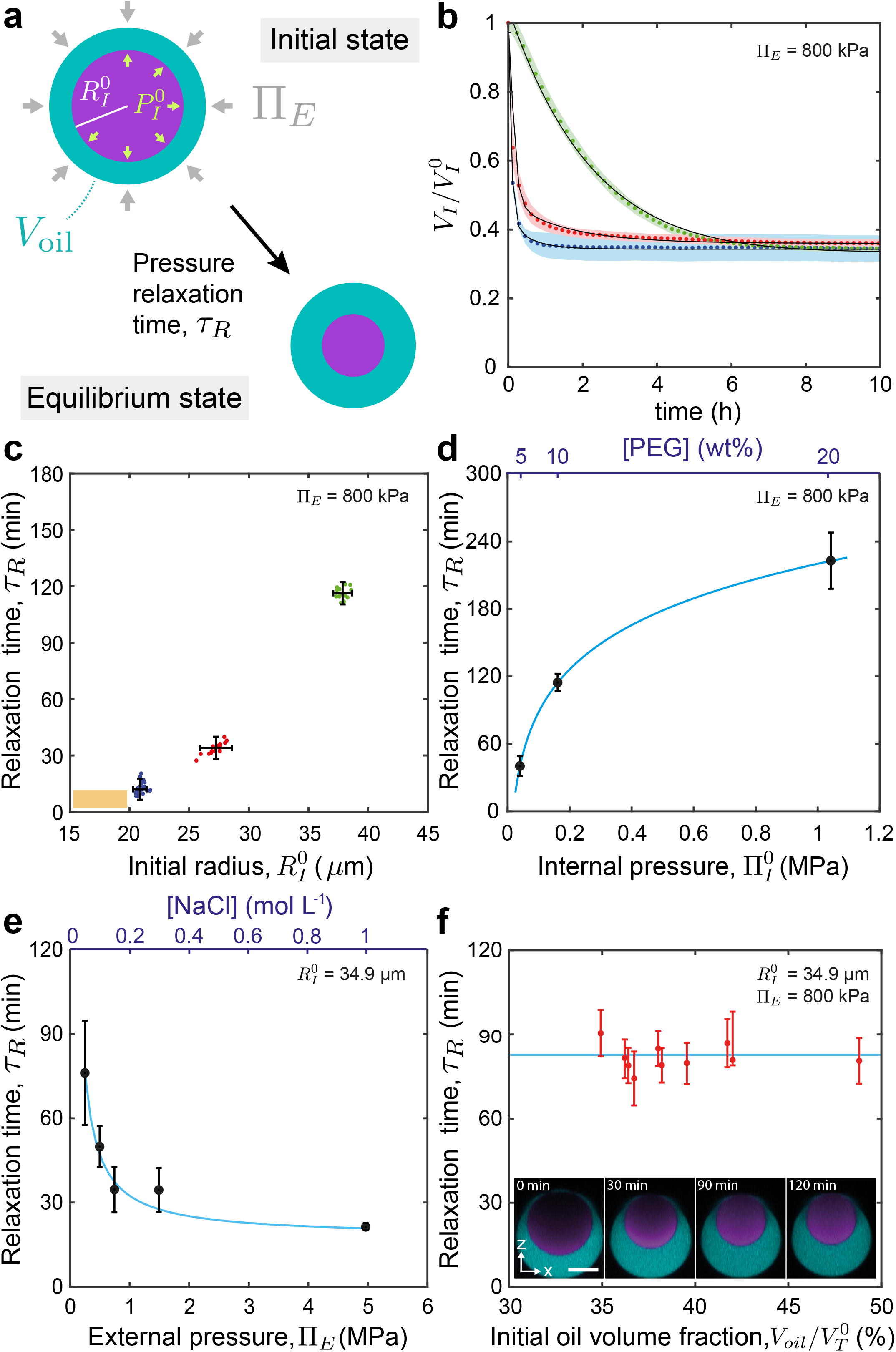
Pressure equilibration timescales of double-emulsion droplets. **a**, Sketch showing a double-emulsion droplet of initial inner pressure 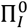 and volume 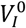 (or radius 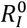) and initial oil volume 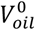, reducing its volume to the equilibrium values over a timescale *τ_R_*. **b**, Inner droplet volume relaxation (normalized to the initial volume 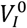) for double-emulsion droplets of different initial sizes: 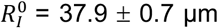, green; 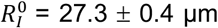, red; 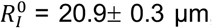, blue (initial PEG concentration (5% w/w) and fixed Π_*E*_). Black lines are exponential fits to the data (Methods). **c-f**, Dependence of the measured equilibrium relaxation timescale (Methods) on the initial inner droplet size, 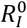 (**c**; initial PEG concentration (5% w/w) and fixed Π_*E*_; N=47 (green), 20 (red), 20 (blue)), the initial internal pressure, 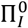 (**d**; fixed 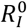 and Π_*E*_, N=13 (5% w/w),18 (10% w/w), and 20 (20% w/w)), the externally imposed osmolarity, Π_*E*_ (**e**; initial PEG concentration (5% w/w) and fixed 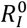, N=11 (0.25 MPa), 15 (0.5 MPa), 12 (0.75 MPa), 14 (1.5 MPa) and 16 (5.0 MPa)), and the initial oil volume fraction, 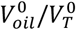 (**f**; initial PEG concentration (10% w/w), fixed 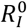 and Π_*E*_), with 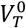 being the initial total droplet volume. Inset in (**f**) shows z-x imaging plane of a droplet relaxing to the equilibrium state (fluorocarbon oil, cyan; fluorescent PEG, purple). Continuous blue lines in **d** and **e** are fits to the data with the form: *y* = *ax^b^* + *c*. Scale bar, 25 μm. All error bars are SD. The orange rectangle in **c** indicates the range of droplet radii used for measurements in living embryos (Fig. 4).

## *In situ* and *in vivo* measurements of osmotic pressure in zebrafish blastomeres

After calibrating double-emulsion droplets, we performed proof-of-principle experiments to measure the osmotic pressure inside blastomeres (cells) of developing zebrafish embryos (Fig. 4a,b). A single double-emulsion droplet was microinjected into the only cell in the embryo at the 1-cell stage (Methods; Fig. 4c), as previously established^9,30,31^. To measure the local value of the osmotic pressure we monitored the volume of the inner droplet over time for over 3 hours, from the 4-cell stage until the cell size became approximately twice the droplet size (Fig. 4c,d). The measured intracellular osmotic pressure values were of 280 mOsm/kg (0.7 MPa) on average (Fig. 4j) and remained largely constant throughout the measurement period (Fig. 4d). These values were similar to those estimated from osmotic shocks *in vitro* for the intracellular osmolarity of cells in culture conditions^18^ (280-300 mOsm). The measured intracellular osmotic pressure should change to the osmotic pressure of the external medium (E3 buffer; Methods) upon dissolution of cell membranes, since the double-emulsion droplet would progressively be exposed to the E3 medium (Fig. 4j; Methods). We used 2% (w/w) sodium dodecyl sulfate (SDS) to dissolve cells’ membranes and completely disperse their contents in the external medium (Fig. 4e; Methods). The osmotic pressure was monitored during the process and found to progressively decrease from its measured intracellular value to the osmotic pressure of the external embryo medium in the presence of 2% w/w SDS (Fig. 4f,j), which was found to be approximately 5-fold smaller than the intracellular osmotic pressure. These results indicate that double-emulsion droplets accurately measure the local osmotic pressure, and that cells (blastomeres) in early embryos tightly regulate their intracellular osmotic pressure through division cycles (cleavages).

**Figure 4.**
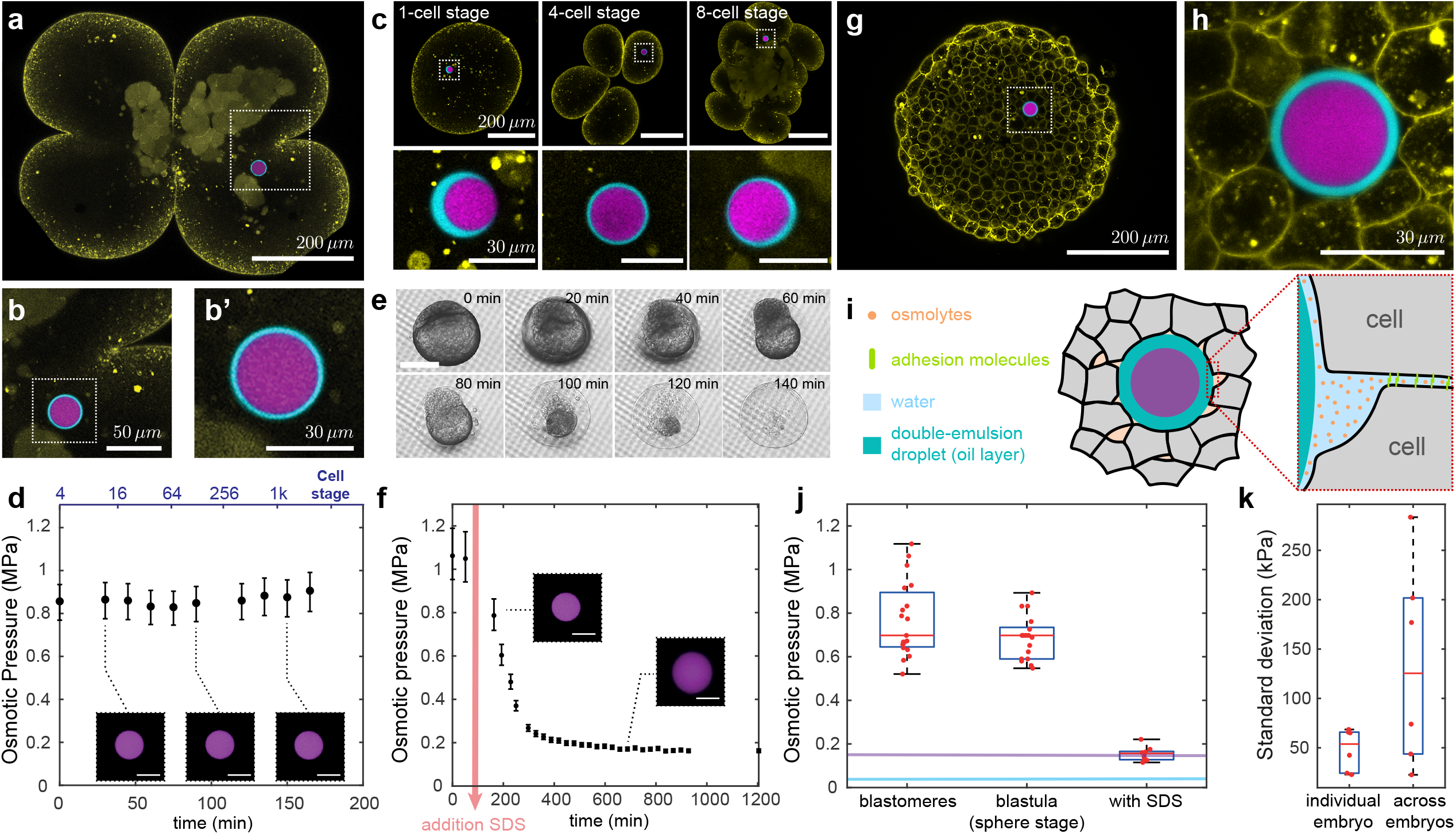
In vivo and in situ measurements of osmotic pressure in blastomeres and in the interstitial fluid of zebrafish embryos. **a**, Confocal section of a *Tg(actb2: mem-NeonGreen)^hm37^* zebrafish embryo transitioning from the 2- to 4-cell stages (membranes, yellow) with a doubleemulsion droplet (fluorescent PEG in inner droplet, purple; fluorocarbon oil, cyan) located in one of the blastomeres (cells). **b-b’**, Close ups of the double emulsion droplet in **a**. **c**, Confocal images of a double-emulsion droplet inside a cell of a developing zebrafish embryo at different developmental stages. Close ups of the double emulsion droplet at each stage. Scale bars, 200 μm (top panels) and 30 μm (bottom panels). **d**, Measured time evolution of the intracellular osmotic pressure in a developing zebrafish embryo. The pressure values were obtained from the calibration curve (Eq. 1; Fig. 2g) after measurements of the inner droplet size at each time point. Inset shows equatorial confocal sections of the inner droplet at different time points. Scale bar, 20 μm. Error bars indicate measurement error for a single droplet (Methods). **e**, Timelapse of a zebrafish embryo in 2% w/w SDS solution imaged in an inverted microscope (transmitted light) and sustained on a porous membrane (Methods). Scale bar, 300 μm. **f**, Measured time evolution of the osmotic pressure during SDS treatment (2% w/w SDS). Insets show inner droplet equatorial confocal sections at different timepoints. Scale bars, 20 μm. Error bars are measurement error for a single droplet (Methods). **g**, Confocal section of a zebrafish embryo blastula at sphere stage (4 hpf; same color code as in **a**) with a droplet inserted in the interstitial fluid between the cells. Scale bar, 200 μm. **h**, Close up showing the equatorial confocal section of the droplet in **g**. Scale bar, 20 μm. **i**, Schematic representation of the droplet in between adhering cells and the presence of osmolytes in the interstitial fluid. **j**, Measured osmotic pressure inside blastomeres, between the cells (interstitial fluid) of the zebrafish blastula (sphere stage) and after SDS treatment. N = 19, 21, 10, respectively. Osmotic pressure of E3 buffer (embryo medium) with (violet line) and without (blue line) 2% w/w SDS, measured with a commercial osmometer (Methods). **k**, Measured osmotic pressure variation (standard deviation) of temporal readings in individual embryos and across the different embryos (Methods). N=6.

## Osmotic pressure of interstitial fluid in developing zebrafish embryos

Beyond intracellular osmotic pressure, the osmotic pressure of the extracellular interstitial fluid located between the cells (Fig. 4i) has also been shown to play an important role in morphogenetic events. To measure the osmotic pressure of the interstitial fluid, we injected a single double-emulsion droplet in between the cells of the zebrafish embryo blastula at sphere stage (Fig. 4g,h; Methods) and monitored the volume of the inner droplet (Methods). The measured osmotic pressure of the interstitial fluid was found to be approximately 0.7 MPa, corresponding to an osmolality of 280 mOsm/kg on average, nearly identical (within error) to the measured intracellular value of osmotic pressure in blastomeres (Fig. 4j). This value is similar to the average osmolality of 260 mOsm/kg measured in whole zebrafish blastula explants^34^, showing that our *in vivo* and *in situ* readings are consistent with these previous *ex vivo* observations. While we could not measure the intracellular osmotic pressure of cells of the blastula at sphere stage because of their small size, our measurements suggest that the intracellular osmotic pressure and the osmotic pressure of the interstitial fluid are constantly balanced, as a mismatch in such high osmotic pressures could be fatal for cells. In contrast, the pressure of the embryo external medium is 17-fold lower than both the interstitial fluid pressure and the intracellular pressure (Fig. 4j).

## Discussion

This work shows that double emulsion droplets can be used as non-invasive, precise and robust osmotic pressure sensors to locally measure osmotic pressure *in vivo* and *in situ* within 3D multicellular systems, such as developing embryos, both intra- and extra-cellularly. Using calibrated double-emulsion droplets, we quantified the osmotic pressure inside cells as well as in the interstitial fluid between the cells of living zebrafish embryos.

The measured values of osmotic pressure reported herein are in agreement with previous *in vitro* inferences from osmotic perturbations of animal cells in culture conditions^17,18^, and are similar to those values estimated from plasmolysis in plant or fungal cells^29^. A previous measurement of the interstitial fluid osmolarity in zebrafish tissue (blastula) explants^34^ reported similar values to those obtained in our measurements. However, those experiments required the destruction of the sample and could only obtain an average value of osmolarity for the entire tissue explant. Our local, *in situ* measurements show that intracellular osmotic pressure is constant throughout the first divisions in zebrafish embryos, and that osmotic pressures inside cells and in the extracellular spaces (interstitial fluid) are balanced. However, we find that the osmotic pressures in the embryo are 17-fold higher than that of the medium external to the embryo, showing that embryos are able to maintain a large osmotic pressure difference between their interior and the surrounding environment. It is possible that the vitelline membrane surrounding the embryo mechanically supports this large pressure difference, similarly to cell walls in bacterial or plant cells, which feature similar intracellular osmotic pressures as those reported here for cells of zebrafish embryos. These results suggest that osmolarity is highly regulated both in cells and in the interstitial fluid, in agreement with our observations showing a much smaller variability in osmotic pressure within a given embryo than across different embryos (Fig. 4k).

Previous *in vitro* studies have shown that several cell behaviors, from cell migration^26,48^ to cortical cell mechanics^17^ as well as cell and nuclear size^13^, depend on the cell’s osmotic pressure. Other works have shown the existence of spatial gradients in extracellular spaces that lead to gradients in tissue stiffness during posterior body axis elongation in zebrafish^9^. Gradients in extracellular spaces can strongly affect tissue hydraulics^23,25,49,50^, which depends on the local regulation of interstitial fluid osmolarity. Finally, the formation of the blastocoel and other embryo cavities, as well as the formation of lumen during organogenesis depend directly on the control of osmotic pressure in these structures^19,24,27,49^. The ability to locally measure osmotic pressure in 3D multicellular systems opens new avenues to study its role in all these cellular and developmental processes, during both embryogenesis and in disease states.

## Data Availability Statement

The data that supports these findings are available upon request.

## Code Availability Statement

The custom-made image analysis code used in this work is available upon request.

## Acknowledgements

We thank I. Lim (UCLA) for sharing custom fluorinated dyes, S. Megason (Harvard Medical School) for providing *Tg(actb2:mem-NeonGreen)^hm37^* embryos, A. Kickuth (Brugués Lab, MPI CBG, Dresden) for sharing her mounting method of early zebrafish embryos and K. Mentoani for SDS screening of zebrafish embryos. We also thank all members of the Campàs lab (especially Carlos Gomez) for discussions and technical support, the CNSI microfluidics and cleanroom facilities at UCSB as well as the UCSB Animal Research Center for support. AV was supported by a postdoctoral fellowship from the Swiss National Foundation (P400PB_191065). This work was supported by the National Institute of General Medical Sciences of the National Institutes of Health (R01GM135380 to ES and OC), and the Deutsche Forschungsgemeinschaft (DFG, German Research Foundation) under Germany’s Excellence Strategy – EXC 2068 – 390729961– Cluster of Excellence Physics of Life of TU Dresden.

## Author Contributions

AV and OC designed research; AV, MP, S-TY performed experiments; AV and SK analyzed the data; SK developed image analysis codes; JP and YL provided technical assistance with microfluidics; ES provided technical expertise; AV and OC wrote the paper; OC supervised the project.

## Online Methods

### Microfluidic device fabrication

The microfluidic devices for producing double emulsions were made of poly(dimethyl siloxane) (PDMS Sylgard 184, Sigma Aldrich, Cat# 761036) and fabricated using soft lithography^51,52^. The template for the double emulsion droplet microfluidic device is based on an existing design^43^. The dimensions of the device were adjusted to achieve the desired droplet sizes. Specifically, we used two different flow focusing devices with different dimensions of the main channel, namely 100 μm width and 60 μm height for the large one and 30 μm width and 30 μm height for the small one. The size of the droplets generated depended on the channel geometry^53,54^. Surface activation of the PDMS devices was done with plasma treatment (Plasma Harrick PDC-32G). Then, a solution containing a cationic polymer, 2% w/w pollydiallyldimethylammonium chloride (PDADMAC, Sigma Aldrich, Cat# 409014) and 1M NaCl (Sigma, Cat# S9888), was used to render the main channel downstream of the 3D junction hydrophilic. A solution of 2% v/v trichloro(1H,1H,2H,2H-perfluorooctyl)silane (Sigma-Aldrich, Cat# 448931) was used to obtain fluorophilic injection channel upstream of the 3D junction.

### Double emulsion droplet composition

The inner droplet was composed of an aqueous solution containing either 5%, 10% or 20% w/w poly(ethylene glycol) (PEG, Sigma, *M_w_* = 6 kDa, Cat# 81260), corresponding to osmolalities of 16mOsm/kg, 65.33 mOsm/kg and 420mOsm/kg (Fig. S3), respectively, and 0.01% w/w of mPEG-Rhodamine (*Creative PEG Works*, Mw = 5 kDa; Cat# PSB-2264) or mPEG-Fluorescein (*Creative PEG Works*, Mw = 5 kDa; Cat# PSB-2254). The concentration of PEG in the inner droplet defines the internal osmotic pressure of the droplet, as PEG cannot go through the oil layer (Extended Figure 1). The presence of PEG also facilitates the generation of double emulsion droplets because it increases the solution viscosity. mPEG-Rhodamine was added at a much smaller concentration to enable droplet size measurements using confocal microscopy. The oil layer surrounding the inner droplet was composed of a fluorinated oil, namely hydrofluoroether (HFE) Novec™ 7700 (3M; ID 7100094084), containing a fluorinated surfactant Krytox-PEG (RAN BioTech Cat# 008) at a 2% w/w concentration, which is a triblock surfactant that has two perfluorinated blocks that are separated by a PEG-based block^55,56^. For imaging purposes, 0.025 mM of custom-made fluorinated dye F_86_Cy5^46^ was added to the oil phase.

### Production of water-in-oil-in-water double emulsion droplets

Each phase was injected in the flow focusing microfluidic device^43^, with each flow rate (Fig. 2b; inner flow rate, *Q_i_*, magenta; oil flow rate, *Q_m_*, cyan; outer flow rate, *Q_o_*, blue) being independently controlled by a different syringe pump (New Era Pump System Model #1000). In addition to the phases described above for the inner droplet and the oil layer, the external aqueous phase to generate the droplet contained 10% w/w partially hydrolyzed poly(vinyl alcohol) (PVA, Sigma, *M_w_* = 13-23 kDa, Cat# 363170). Water-in-oil-in-water double emulsion drops with diameters ranging from 25 to 120 μm were formed using the two flow focusing devices described above. Control over the general size of the droplet was achieved by two parameters: the type of device and outer flow rate. For droplets with initial diameter ranging from 60 to 120 μm, we used the large device and outer flow rate *Q_o_* ranging between 3000 to 6000 μL/h. For droplets with initial radius between 20 to 60 μm, we used the small device and outer flow rate *Q_o_* ranging between 300 to 1700 μL/h. To change the initial oil volume fraction, 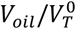, we used a large device and kept *Q_o_* constant at 4500 μL/h, while the ratio *Q_i_/Q_m_* was adapted from 1:1 to 8:3.

### Osmotic pressure calibration

The osmolality of all aqueous solutions used for calibration and testing was measured with an osmometer (Advanced instruments, Model 210, Case # 34458). Conversion from osmolality, *π_osm_*, to osmotic pressure, Π, was done using the Van’t Hoff Law for dilute solutions, namely Π = *π_osm_RT*, with *R* the gas constant and *T* the temperature in Kelvin^17,18^.

In order to relate the internal volume of the droplet to the external pressure we produced droplets with initial inner radius of 33.5 ± 0.6 μm. Those droplets were subsequently placed into NaCl solutions with calibrated concentrations of 0.05 M, 0.1 M, 0.15 M, 0.3 M, 0.5 M and 1 M. For ionic and dilute solutions, the osmotic pressure is related to the concentration as follow. The osmolality *π_osm_* = *nϕc*, with *n* being the number of particles in which the compound dissociates, *ϕ* being the degree of dissociation of the solute and c the solute concentration. In the case if NaCl, *n =* 2 and *ϕ* = 1. Knowing *n, ϕ*, and c, for the NaCl solution, we obtained *π_osm_* and the osmotic pressure according to Π = *π_osm_RT*. This provided a solution of well-known osmotic pressure that was used to calibrate the droplets.

### Storage and osmotic pressure calibration

All droplets produced with microfluidic devices were generated and initially stored in 10% w/w PVA aqueous solution with osmolality of 79 mOsm/kg (or 200 kPa in osmotic pressure).

The osmolality of cell culture media measured in the osmometer was 839 kPa or 338 mOsm/kg. E3 embryo media was composed of NaCl (290 mg/L), KCl (13.33 mg/L), CaCl_2_ (4.83 mg/L), MgCl_2_ (81.5 mg/L) and methylene blue (1 vol%, 100μL/L). The measured osmolality of the E3 embryo media was 11 mOsm/kg (27.3 kPa), which increased to 48 mOsm/kg (118 kPa) when SDS was added at a 2% w/w concentration.

The cell culture media used in calibration experiments was composed of RPMI 1640 (ThermoFisher, Cat# 11875093), supplemented with 1% w/w Pencillin-Streptomycin (ThermoFisher, Cat# 15140122) and 10% w/w Heat Inactivated Fetal Bovine Serum (ThermoFisher, Cat# MT35011CV).

### Characteristic relaxation time

In order to characterize the characteristic timescale of pressure equilibration, we monitor the droplet volume changes over time and fit an exponential decay to the data. The characteristic relaxation timescale is the timescale of the exponential fit.

### Zebrafish husbandry and fish lines

Zebrafish (*Danio rerio*) were raised and bred as described previously^57^. Animals were raised and experiments were performed following all ethical regulations and the protocols approved by the Institutional Animal Care and Use Committee (IACUC) at the University of California, Santa Barbara. A *Tg(actb2:mem-NeonGreen)^hm37^* transgenic line was used for ubiquitous labeling of cell membrane of zebrafish embryos.

### Injection of double emulsion droplets in zebrafish embryos

Zebrafish embryos at 1-cell stage were chemically dechorionated by 1 mg/mL of pronase (Roche, Cat# 10165921001) in E3 buffer. Embryos at sphere stage were dechorionated manually. Embryos were all microinjected with double emulsion droplet in 0.1 M KCl (Sigma, Cat# P3911) solution using a picolitre injector (Warner Instruments LLC, PLI-100A). Micropipettes for microinjection were made from microcapillaries (World Precision Instrument; TW100F-4) using a Sutter P-1000 needle puller and were coated with 2% w/w PDADMAC in 1M NaCl to avoid rupture of the double emulsion droplet inside the micropipette. The diameter of the inner droplets of the double emulsions ranged between 20 – 35 μm. Double emulsion droplets were back-loaded into the microneedle, which tends to accumulate at the tip of the needle due to gravity. Injection pressure was tuned to achieve the injection of single droplets in the embryo.

### Mounting and imaging of double emulsion droplets in zebrafish embryos

All images were acquired using a Zeiss LSM710 laser scanning confocal microscope. Imaging of zebrafish embryos injected with double emulsion droplets were mounted in 0.75% low-melting point agarose (Invitrogen; Cat# 16520-050) mixed with 25% OptiPrep™ density gradient medium (Sigma; Cat# D1556) in E3 buffer (without methylene blue)^58^ in a glass-bottom dish (MatTek; P35G-1.5-14-C) with two layers of silicone isolators (Electron Microscope Sciences, Cat.# 70336-61). For SDS treatment, a 40-μm nylon mesh, which was cut out from cell strainers (Fisher Scientific; Cat# 22363547), was used to provide a porous seal instead of a coverslide. SDS treatment was administered at 128-cell stage of the zebrafish development and measurement of the drop size was manually performed every 15 min for 4h.

Images of early development zebrafish were taken using a 10x air objective (EC Epiplan-Neofluar 10× 0.25, Carl Zeiss Inc.). For measurements of volume changes in double emulsion droplets, 3D timelapses of droplets were acquired using a 40X water immersion objective (LD C-Apochromat 1.1W, Carl Zeiss Inc.) at 25 °C. Confocal section in z were between 5-10 μm with the 10X objective and 1-2 μm for the 40x objective.

### Imaging of double emulsion droplets for calibration purposes

3D confocal timelapses of double emulsion droplets were acquired on a Zeiss LSM710 laser scanning confocal microscope with a 40x water immersion objective (LD C-Apochromat 1.1W, Carl Zeiss Inc.) at room temperature.

### Analysis of inner and outer droplet volumes

To quantitatively obtain the droplet’s size from imaging data, we developed a custom-made Matlab code. First, maximum intensity projections (MIP) of the measured z-stack of multiple droplets in a region of interest were obtained for the inner droplet at every timepoint. We focused on the inner droplet because changes in outer droplet volume follow changes in inner droplet volume (Fig. 2e). Individual timelapses of inner droplets’ MIPs were segmented by thresholding a grayscale image with an input threshold value. Segmentation artifacts smaller than a critical object size are removed and binary erosion operation and binary dilation operation are applied consecutively to generate smooth droplet interfaces. Individual droplets were then labelled at each time point and tracked over time based on the shortest distance criterion between consecutive time points. For each droplet identified in the segmented image, the droplet area *A* was computed by counting number of pixels. Since the inner droplets maintained spherical shape, the inner droplet volume *V_I_* was obtained from

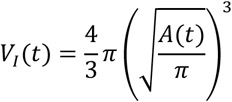

